# A Hybrid Method for Ultrasound-Based Tracking of Skeletal Muscle Architecture

**DOI:** 10.1101/2022.04.20.488774

**Authors:** Jasper Verheul, Sang-Hoon Yeo

## Abstract

Tracking skeletal muscle architecture using B-mode ultrasound is a widely used method in the field of human movement science and biomechanics. Sequential methods based on optical flow algorithms allow for smooth and coherent muscle tracking but are known to drift over time. Non-sequential feature detection methods on the other hand, do not suffer from drift, but are limited to tracking only lower-dimensional features. They are also known to be sensitive to image noise, and therefore often result in highly irregular tracking patterns. Building on the complimentary nature of both approaches, we present a novel fully automated hybrid muscle tracking approach that combines a sequential feature-point tracking method and a non-sequential method based on Hough transform. Tibialis anterior fascicle pennation angle and length, and central aponeurosis displacement, were measured in five healthy individuals during isometric contractions at five different ankle angles. Our hybrid method was demonstrated to significantly (p < 0.001) reduce drift compared to two sequential methods, and curve irregularity was significantly (p < 0.001) decreased compared to a non-sequential method. These findings suggest that the proposed hybrid approach can uniquely mitigate drift and irregularity limitation of individual methods used for tracking skeletal muscle architecture. Fully automated muscle tracking allows for convenient analysis of large datasets, whereas automatic drift correction opens the door for tracking muscle architecture in long ultrasound recordings during common movements, such as walking, running, and jumping without the need for manual intervention.

## 1 Introduction

Ultrasonography has been used in a range of medical fields to non-invasively examine the cardiovascular and musculoskeletal systems in vivo. In the field of human movement science and biomechanics, brightness-mode (B-mode) ultrasound is primarily used to evaluate the anatomical, architectural, and mechanical properties of skeletal muscles and tendons [1]. Examples of properties that are extracted from ultrasonographic images include the thickness and cross-sectional area of muscles and tendons, the pennation angle and length of muscle fascicles, the stiffness of tendons and aponeuroses, and the stresses and strains in tendinous tissues. Extracting muscle-tendon properties from ultrasound images is, however, not straightforward and relies heavily on postprocessing procedures.

Seminal work in the area of muscle ultrasonography has relied primarily on the manual annotation and analysis of muscle properties [2], [3]. However, this process is subject to experimenter bias and human error and, more importantly, can be laborious when processing large datasets and/or long image sequences (i.e., videos). Various (semi-)automated methods have, therefore, been suggested to allow for objectively detecting and tracking muscle architectural properties from ultrasound images and videos. These methods can be divided into two main groups: sequential and non-sequential methods.

Sequential methods are typically based on optical flow estimation algorithms. These algorithms estimate the displacement of feature points, i.e., points located on a specific feature in an image, from one image frame to the next. By summating these frame-to-frame displacements over time, feature-point trajectories can be obtained. Subsequently, the displacement or deformation of a region of interest (ROI), such as a muscle fascicle or aponeurosis, can be estimated by tracking a group of feature points defined in that region. Since users can flexibly define such ROIs and the tracked feature points provide direct two-dimensional information of local displacements, these methods can be widely applied to different scenarios of ultrasound-based muscle tracking [4]–[7]. Importantly, sequential tracking methods are known to be robust against image noise, and therefore allow for spatiotemporally smooth and coherent tracking.

Nevertheless, like other differential-based methods, a serious drawback of sequential tracking is drift, which is the accumulation of small displacement-estimation errors leading to the ROI moving away from its actual location over time (see e.g., [5]). Drift can substantially reduce the measurement accuracy of muscle properties over time, especially for longer ultrasound recordings, rapid muscle contractions, and low image sampling frequencies or resolutions.

Solutions for correcting drift in sequential muscle tracking methods are yet scarce. Farris and Lichtwark have suggested a ‘key-frame correction’ algorithm [5]. Key-frame correction assumes that muscle architecture is identical at recurring reference states throughout a movement, such as the ground contact phase during walking. These reference states can either be manually selected time instances or identified from additional data (e.g., ground reaction forces). However, this algorithm relies on the assumption that movements are periodic, which is often not the case. Moreover, criteria for manual adjustments can be difficult to determine, and additional information is not always available (e.g., for walking or running on a non-instrumented treadmill). It is, therefore, preferable that drift-correction methods do not rely on any manual interventions or additional data besides the ultrasound images under analysis.

As an alternative to sequential approaches, single-image-based, non-sequential methods have been used to detect and track muscle architecture. Non-sequential methods typically detect features, such as lines or specific patterns, in individual images without considering the sequential relationship between neighboring images. Examples of such approaches are the Hough transform [8]–[10] and the Radon transform [11], [12]. Muscle properties that can be extracted using these methods include the fascicle pennation angle and length, and muscle thickness, which can be arrayed for multiple images to obtain changes in muscle architecture over time. Since each image in a sequence is analyzed separately, non-sequential methods have the major advantage of not suffering from drift, but also have several shortcomings. First, not considering the temporal coherence of neighboring images makes non-sequential methods sensitive to image noise, especially the speckle noise typical for ultrasonography. Second, detected features can be easily distorted by irregularities that are often observed in ultrasonographic muscle images (e.g., connective tissues or blood vessels). This can lead to substantially more irregular results compared to sequential methods. Third, whereas sequential methods can flexibly extract various features of deformation, non-sequential methods are limited to individual features. For example, line detectors (e.g., Hough or Radon transform) are limited to estimating the slope of detected line segments (e.g., fascicle pennation angles or aponeurosis lines), leaving image features that do not involve line orientations undetectable.

Building on the complementary nature of sequential and non-sequential tracking methods, this paper presents a hybrid method for ultrasound-based tracking of skeletal muscle architecture. We describe a novel approach that combines two existing tracking methods that have previously been used in the field: 1) sequential feature-point tracking using a Kanade-Lucas-Tomasi algorithm and 2) non-sequential tracking using Hough transform. This novel hybrid method is then applied for tracking the human tibialis anterior muscle during isometric contractions. The key aims of our approach are to:

1. Quantify regular (i.e., smooth) time-series waveforms for fascicle pennation angle and length, and aponeurosis displacement.
2. Eliminate drift without the need for data other than ultrasound or manual input, while maintaining temporal coherence.

## 2 Methods

### 2.1 Data collection

Five healthy individuals (four males, one female, age: 25±4 yrs) agreed to participate in this study. Participants were well trained, free of lower-limb injuries for at least six months, and had no history of neurological or cognitive impairment. A full description of the experimental protocol was provided to all participants to which they agreed by providing informed consent. This study was approved by the University of Birmingham research ethics committee.Five healthy individuals (four males, one female, age: 25±4 yrs) agreed to participate in this study. Participants were well trained, had been free of lower-limb injuries for at least six months, and had no history of neurological or cognitive impairment. A full description of the experimental protocol was provided to all participants to which they agreed by providing informed consent. This study was approved by the University of Birmingham research ethics committee.

Participants were seated on the chair of a dynamometer (Biodex 3 system, Biodex Medical Systems) with their right knee fully extended and the right foot securely fixed to a footplate. The lateral malleolus was aligned with the center of rotation of the dynamometer arm, with the ankle joint in a 0° neutral position. A linear ultrasound transducer probe (12L-RS probe, Vivid iq system, General Electric Healthcare, probe width: 38 mm, scan depth: 35 mm, frequency: 13 MHz, resolution: 1280×720 pixels) was used to record tibialis anterior muscle architecture. Ultrasound transmission gel was applied to the probe, after which the probe was placed over the belly of the tibialis anterior so that the central aponeurosis was horizontal in the image and moved primarily in horizontal direction during contraction. The probe was then securely fixed to the lower leg using a purpose-made polystyrene fitting and self-adhesive bandage (Figure 1).

**Figure 1:**
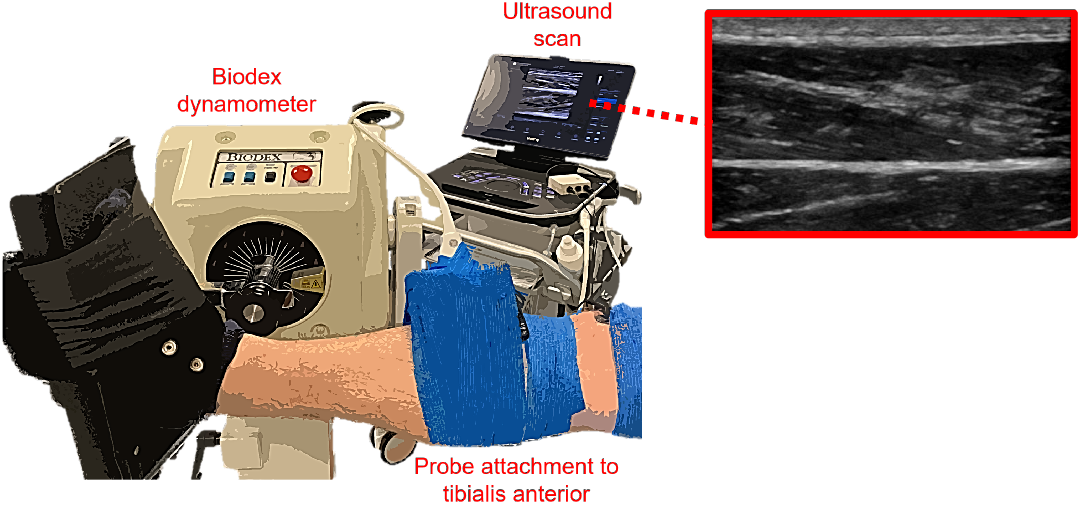
Data collection setup: positioning of the right leg in the dynamometer and ultrasound probe attachment to the tibialis anterior muscle.

Participants first performed a minimum of five unloaded concentric ankle dorsiflexion movements through the whole range of motion in the dynamometer as a warmup, for familiarization purposes, and to check data quality. A series of isometric dorsiflexion trials was then performed at five different ankle angles ranging from 10° dorsiflexion to 30° plantarflexion (i.e., the typical range of motion of the ankle joint during locomotion) in 10° steps and randomized order. At each ankle angle, a total of four twenty-second trials were performed, for which participants were instructed to randomly increase and decrease their isometric dorsiflexion moment at varying speeds and moment levels. The ankle moment profile was visualized on a screen and participants were instructed to change their contraction velocity and/or moment produced during the trials if the moment profile showed little variation, to assure a variety of moment and contraction-velocity levels.

During the trials, tibialis anterior ultrasonography and ankle dorsiflexion moment were recorded synchronously at sampling frequencies of 30 Hz and 1000 Hz respectively. Data were synchronized, stored, and saved via Simulink and MATLAB (version R2021a, The MathWorks). Further processing and tracking of the ultrasound videos were performed in MATLAB.

### 2.2 Hybrid muscle tracking

A hybrid muscle tracking method was developed that combines a sequential (feature-point tracking) and non-sequential (Hough transform) method, that have previously been used separately for muscle tracking purposes [4], [5], [8], [9]. An open-source MATLAB code for the presented hybrid muscle tracking method, including an example trial, is freely available on GitHub (https://github.com/JasperVerheul/hybrid-muscle-tracking).

To start, the location of the superficial and central aponeurosis was identified in each ultrasound image. This aponeurosis-identification process was similar to the approach described by van der Zee and Kuo [8]. Vessel-like structures within each image were computed by filtering the image with a Hessian-based Frangi vesselness filter [13], using an open-access custom MATLAB function [14]. Images were filtered with a Frangi scale range (i.e., Sigma values) between one and two millimeters to allow for detecting wider vessel structures, and binarized using adaptive thresholding. The vessel structures with the largest perimeter in the region from 5-20% (superficial aponeurosis) or 30-60% (central aponeurosis) of the vertical image dimension were selected (Figure 2B). The centroid and orientation of these two remaining vessel structures were then used to define a line representing both aponeuroses (Figure 2C), and their boundaries were cropped to fall within one millimeter (i.e., 14 pixels) of the defined aponeurosis lines.

**Figure 2:**
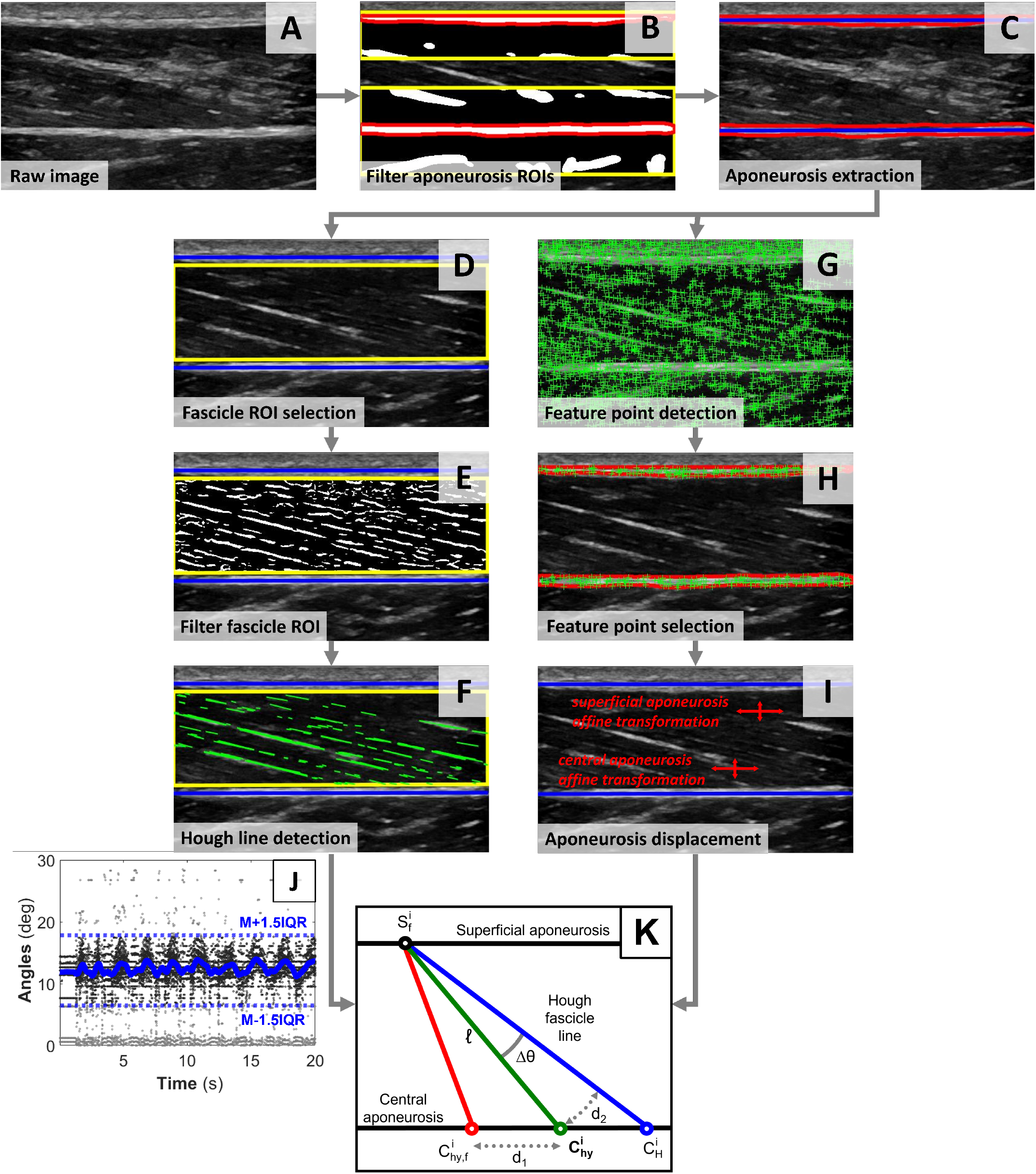
Step-by-step visual representation of the hybrid tracking methodology. A: raw ultrasound image. B: Frangi vesselness filtered aponeurosis regions of interest (yellow box) with the largest vesselness structures (red). C: selected superficial and central aponeuroses boundaries (red) with aponeurosis lines (blue). D: aponeurosis boundaries with fascicle region of interest (yellow box). E: Frangi vesselness filtered fascicle regions of interest (ROI). F: detected Hough lines from Hough transform (green). G: detected feature points (green +). H: selected feature points within fascicle boundaries. I: aponeurosis displacement was based on tracked feature points. J: smoothing spline curve fitted to length-weighed Hough angles in all images. K: optimal hybrid intersection point with the central 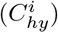 based on distances *d*_1_ and *d*_2_.

Based on the aponeurosis lines, a fascicle region of interest (ROI) was defined for each image. The fascicle ROI was defined as the area between one millimeter below and above the superficial and central aponeurosis lines respectively, over the whole width of the image (Figure 2D). A Hough-transform [15] algorithm was then used to determine the pennation angle within the fascicle ROI. This process involved a series of steps. First, the fascicle ROI was filtered with a Frangi vesselness filter [13], [14] as described in the previous paragraph, with the only difference being the use of a smaller scale range to allow for identifying the finer vessel structures of the muscle fascicles (Figure 2E). Second, the fascicle ROI was stretched vertically to magnify the length of the fascicle lines within the ROI, as Hough transform is known to perform better for longer lines [16]. Since stretching a ROI beyond three times its original dimension can lead to gaps in the magnified lines [16], the fascicle ROI was stretched by a factor three, using bicubic interpolation. Third, the Hough transform was applied to the filtered and stretched fascicle ROI. The rho resolution (i.e., the Hough transform bin spacing on the rho axis of the accumulator) and angle interval (i.e., the stepwise increase over the range of angles of the accumulator) were set to 1 pixel and 0.1° respectively. Hough lines associated with the 50 highest peaks in the accumulator, with a minimal length of 0.5 mm, were selected. Hough lines were discarded if the line angle was: smaller than 0° (i.e., negative pennation angle); 45° due to the Hough transform’s bias towards diagonal lines [17]; or equal to the central aponeurosis orientation to avoid edge effects of the fascicle ROI. The remaining Hough lines were used to determine the pennation angle for each image (Figure 2F).

A Kanade-Lucas-Tomasi feature-tracking algorithm [18], [19] was used to track the displacement and deformation of the superficial and central aponeurosis over time. Feature points were detected as corners within each image using a minimum eigenvalue criterion [20] (Figure 2G), from which the feature points within the aponeurosis boundaries were selected (Figure 2H). Feature points were then tracked on the next image of the video using a point-tracker object, with four pyramid levels, a block size of 31 by 31 pixels, a maximal bidirectional error of one pixel, and a maximum of 50 iterations. Tracked feature points with a confidence score of less than 90% were discarded. Feature points within the aponeurosis boundaries were renewed for each image to assure a maximal number of trackable points. Displacements of the feature points within each of the aponeurosis boundaries were used to determine the affine geometric aponeurosis transformations. However, if less than 20% of all the feature points had been tracked between two images, the transformation was regarded as invalid.

After calculating the Hough-transform angles within the fascicle ROI and affine aponeurosis transformations for each image, a series of postprocessing steps was performed. Hough lines with an angle that was outside one and a half interquartile range from the median of all angles in the video were discarded to remove outliers. A smoothing spline curve was fitted to the remaining Hough-line angles determined for all images in the video (Figure 2J). Angles were weighed according to their line length, and the resulting curve was smoothed with a ten-point moving mean filter. Likewise, a ten-point moving mean filter was applied to the position of the central aponeurosis line endpoints, after which its angle was determined.

As a starting position for hybrid tracking, an initial fascicle line was defined for the first image frame in the video. This line was drawn from the center of the fascicle ROI with an angle equal to the Hough-transform angle in the first image frame. The intersection points between this initial line with the superficial and central aponeurosis lines were defined as 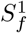 and 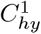 respectively. These intersection points were then updated iteratively. In the *i*^*th*^ image frame, the intersection points in the previous image frame, 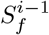 and 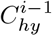 were updated using the affine transformations obtained from feature tracking. The updated intersection point with the superficial aponeurosis was set to 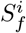, while the intersection point with the central aponeurosis was named 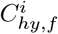. In addition, a Hough-transform-based intersection point 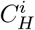 on the central aponeurosis was determined using a newly estimated Hough line drawn from 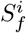 (Figure 2K). Our hybrid algorithm then finds an opti-mal point 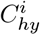 on the central aponeurosis from 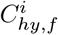 and 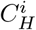 in the following way:

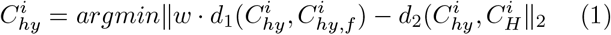

in which *d*_1_ (·) is the length between 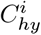 and 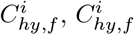 is the feature-tracked point 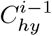 onto the current image frame, *d*_2_ (·) is the arc length defined as Δ*θ* · *l* (see Figure 2K), and *w* is the weight factor. Weight *w* determines the preference for using 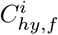 compared to 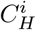 for determining the hybrid point. Since Hough transform generally comes with a higher level of noise compared to feature-point tracking, a reasonably high weight biasing towards feature-point tracking (12.0 in our case) was used for the stability of the iterative optimization procedure. Pseudocode outlining the procedure is presented in Figure 3.

**Figure 3:**
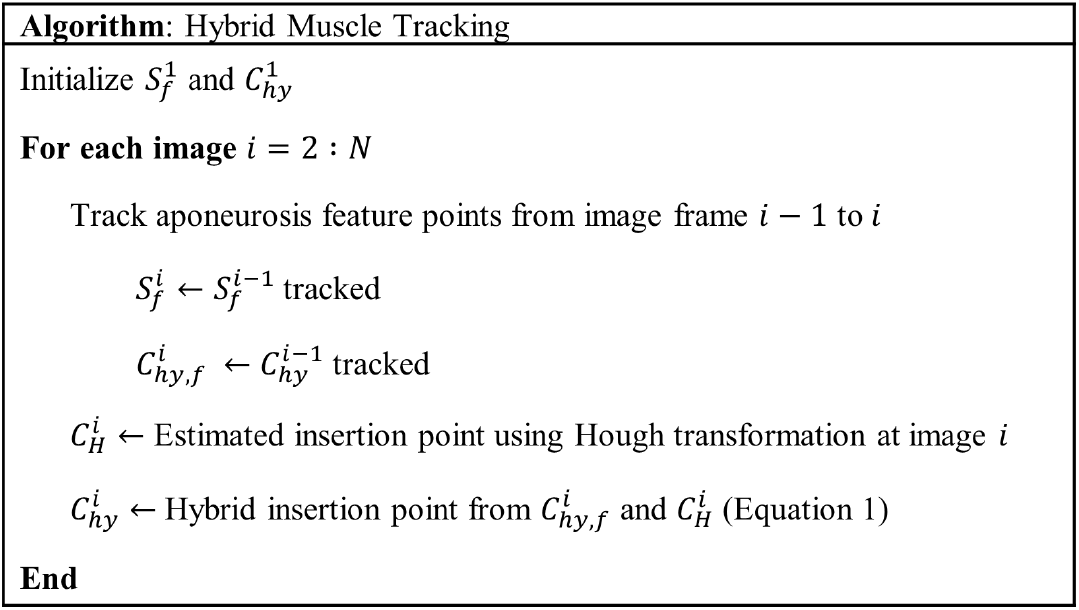
Pseudocode for the hybrid muscle tracking algorithm.

Once 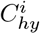 and 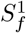were determined, the corresponding fascicle length and pennation angle, and central aponeurosis displacement in that image frame were calculated. Fascicle length was calculated as the distance of the line from 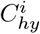 to 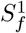. Fascicle pennation angle was defined as the angle between this fascicle line and the central aponeurosis. Aponeurosis displacement was defined as the resultant displacement of 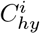 from *C*^0^.

### 2.3 Result analysis

The tracking performance of our hybrid method for fascicle pennation angle was assessed by the amount of drift observed. Since the Hough transform is not subject to drift over time, the pennation angle tracked by the Hough transform was taken as the reference signal for measuring the amount of drift in other methods. With respect to this, drift in pennation angle tracking was then determined for the hybrid method, a feature-point tracking method (i.e., 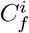 is updated iteratively for each image frame using the affine aponeurosis transformation, similar to 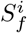 described above), and an existing popular semi-automated tracking method (i.e., UltraTrack version 4.2 [5]). For UltraTrack, ultrasound videos were exported to the required format and the region of interest and an initial fascicle were manually identified on the first image frame of each video, after which the video was processed and the results exported to MATLAB (for more details on this process, see [5]). Since UltraTrack defines pennation angle as the angle of the fascicles relative to the horizontal, pennation angles were not corrected for the central aponeurosis angle when used for comparison purposes. The mean drift per frame for each video was quantified as:

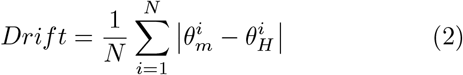

in which Drift is the mean drift per frame (in °) for the pennation angle determined from each tracking method 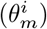 relative to the Hough-transform determined pennation angle 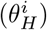 at the *i*^*th*^ image frame of the ultrasound video, and *N* the total number of ultrasound images in each video.

Tracking results for the central aponeurosis displacement were analyzed for irregularity. Small fluctuations in estimated pennation angles can be magnified if displacements are determined from pennation angle only (e.g., Hough-determined angles) and thus provide a highly irregular displacement pattern. Irregularity of the displacements determined from the hybrid, Hough-transform, and UltraTrack methods was, therefore, compared to feature-point tracking. The displacement for UltraTrack was calculated from the fascicle endpoint closest to the central aponeurosis. Displacement irregularity for each method was quantified as:

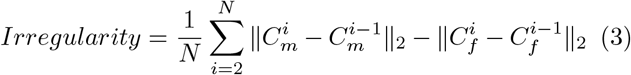

in which Irregularity is the mean irregularity per frame (in mm) of the central aponeurosis displacement profile for each tracking method m relative to the displacements derived from the feature-point tracking method, 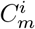 and 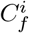are the respective positions of the insertion in the central aponeurosis for each tracking method and uncorrected feature-point tracked point (as described above) at the *i*^*th*^ image, and *N* the total number of ultrasound images in each video.

To compare drift and irregularity between the hybrid, Hough transform, feature-point tracking, and UltraTrack methods, a one-way repeated-measures ANOVA was used. Significance was accepted at p < 0.05. Statistical analysis was performed in SPSS (v.27, IBM, SPSS, Chicago, IL).

## 3 Results

A total of 103 ultrasound videos for the five participants was tracked and processed. Visual screening of the ultrasound videos revealed that nine videos for one participant contained an excessive amount of blur in the images (likely due to improper ultrasound probe attachment to the skin or movement during the experiments). These videos were therefore discarded, leaving a total of 94 twenty-second videos (i.e., 56400 image frames) included in the analysis.

Figure 4 shows two representative example trials which demonstrate that our hybrid tracking method mitigates drift over time. These examples are for trials at 10° ankle plantarflexion (Figure 4, left column) and 10° ankle dorsiflexion (Figure 4, right column) from two different participants. In the first example (Figure 4, left column) the pennation angle determined from the feature point-tracking method suffers from substantial drift over the first six seconds of the trial, which was corrected for by our hybrid method. The mean pennation angle drift this example was 2.72° per frame and 0.3° per frame for the feature-point tracking and our hybrid method respectively. Likewise, the second example (Figure 4, right column) shows substantial drift for the feature-tracking method over the second half of the trial (mean pennation angle drift: 1.14° per frame), but not the hybrid method (mean pennation angle drift: 0.32° per frame).

**Figure 4:**
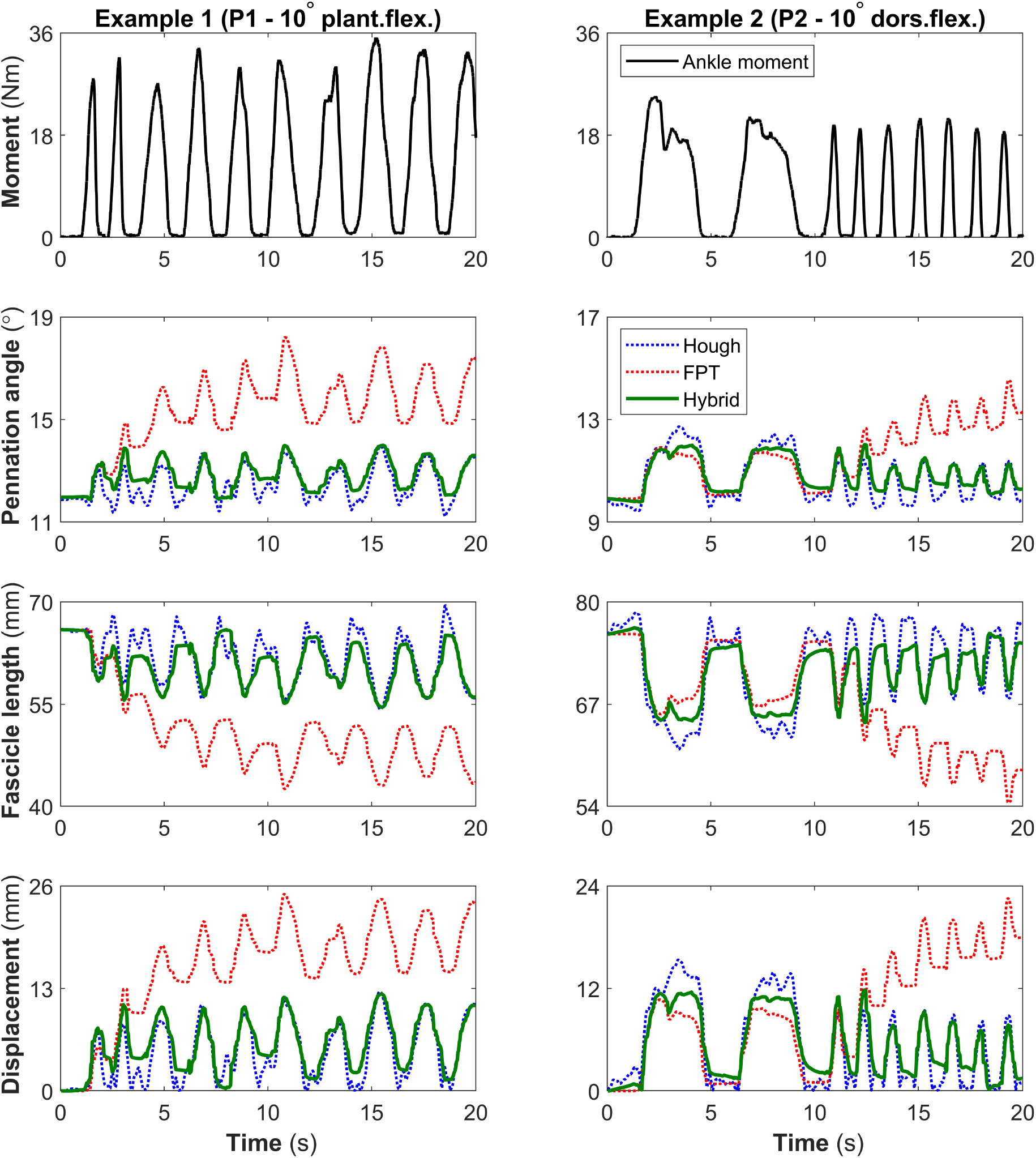
Ankle moment, fascicle pennation angle and length, and central aponeurosis displacement for two representative examples. The hybrid method (green solid line) mitigates drift over time observed in the feature-point tracking method (FPT; red dotted line), using the Hough transform method (blue dotted line). Left: participant one at an ankle angle of 10° plantarflexion. Right: participant two at an ankle angle of 10° dorsiflexion.

Figure 5 shows two representative example trials which demonstrate that our hybrid method can effectively diminish the irregularity of the aponeurosis displacement profile of the Hough-transform method. These examples are for trials at 30° ankle plantarflexion (Figure 5, left column) and 10° ankle plantarflexion (Figure 5, right column) from two different participants. The first example (Figure 5, left column) shows that small variations in Hough-determined pennation angles translate to major fluctuations in the aponeurosis displacement profile. Spikes in the displacement particularly occurred between muscle contractions. In our hybrid method, these additional spikes were substantially mitigated. The mean irregularity of the aponeurosis displacement in the first example trial was 3.13 mm and 0.9 mm per frame for the Hough transform and hybrid method respectively. In the second example (Figure 5, right column) considerable fluctuations in the Hough-determined displacement profile occur primarily during the slower contractions over the second half of the trial. These irregularities are adjusted for in our hybrid method, with mean aponeurosis displacement irregularities of 1.74 mm (Hough) and 0.55 mm (hybrid) per frame.

**Figure 5:**
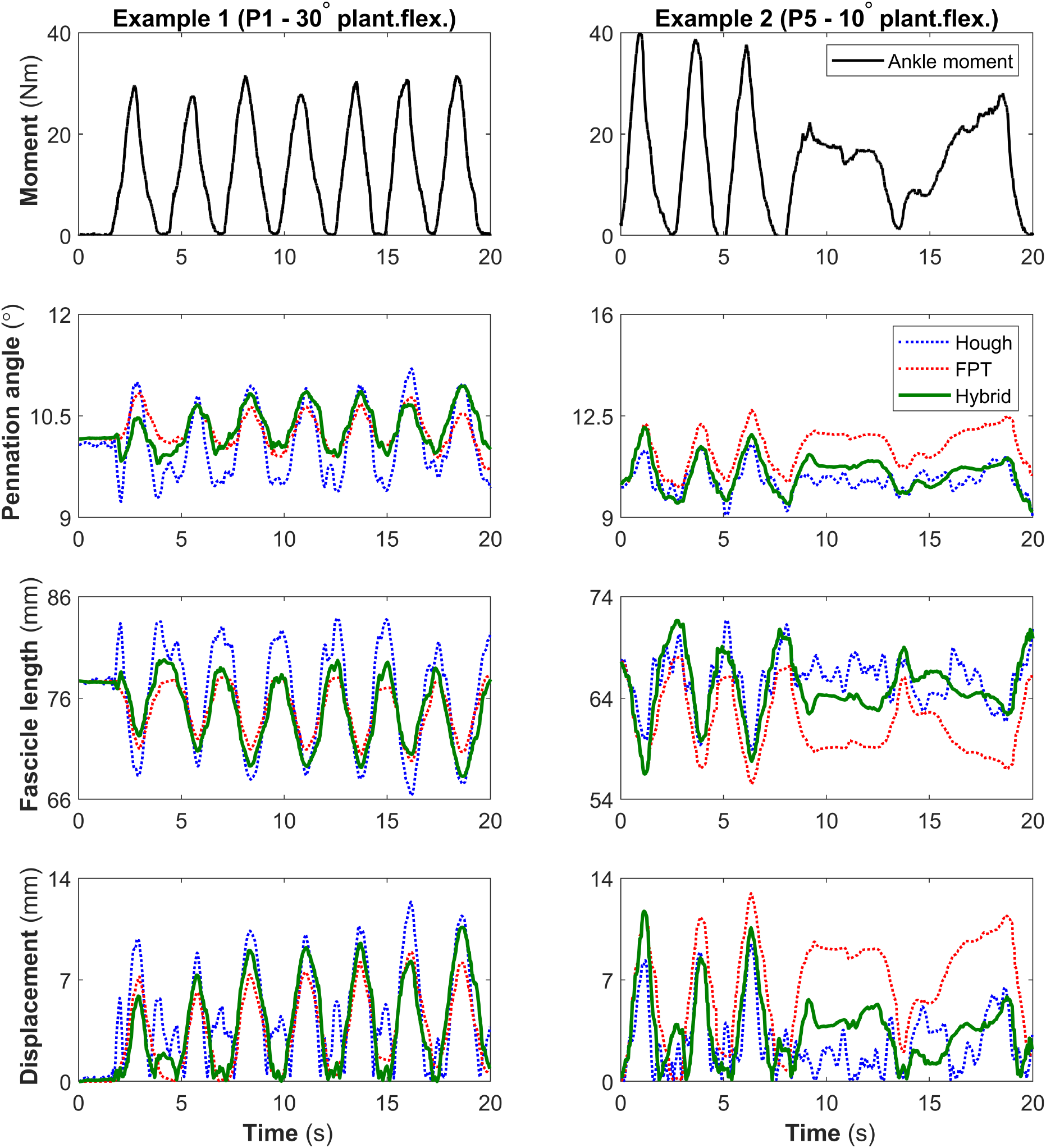
Ankle moment, fascicle pennation angle and length, and central aponeurosis displacement for two representative examples. Our hybrid method (green solid line) reduced aponeurosis displacement irregularity compared to the Hough transform method (blue dotted line), using the feature-point tracking method (FPT; red dotted line). Left: participant one at an ankle angle of 30° plantarflexion. Right: participant five at an ankle angle of 10° plantarflexion.

Figure 6 shows the drift and irregularity results for each trial of all five participants. Mean pennation angle drift across all videos was significantly reduced (p < 0.001) from 1.12±0.8° (UltraTrack) and 0.81±0.63° (feature-point tracking) per frame, to 0.32±0.1° per frame for our hybrid method. Mean irregularity for the hybrid method across all videos was 0.89±1 mm per frame, which was a significant decrease (p < 0.001) compared to the Hough transform (2.8±1.79 mm per frame), but also significantly higher (p < 0.001) than UltraTrack (0.05±0.21 mm per frame).

**Figure 6:**
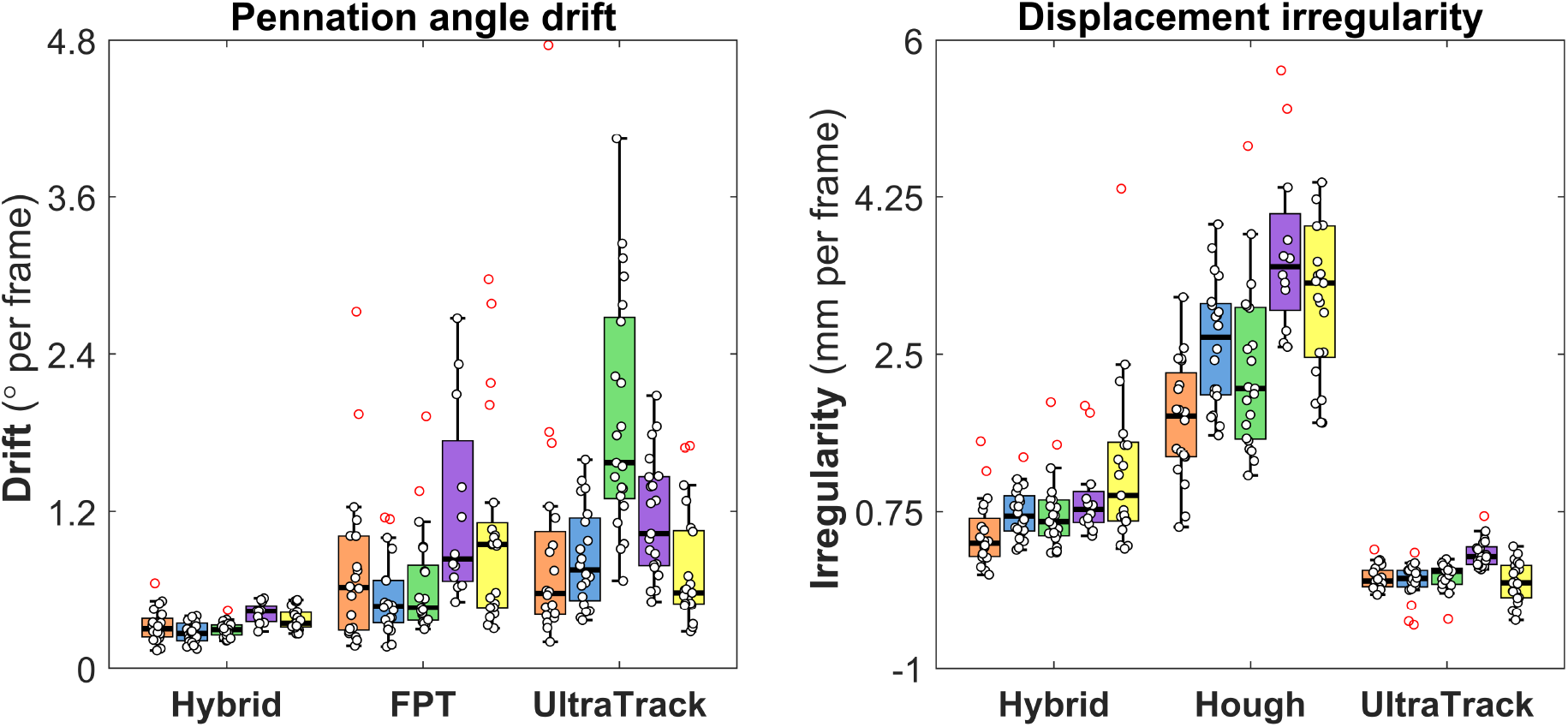
Pennation angle drift (left) and central aponeurosis displacement irregularity (right) boxplot comparisons between the hybrid, Hough transform, feature-point tracking (FPT), and UltraTrack methods. Different colors of the boxplots represent the five participants. Note: to enhance readability, three outliers are not shown in the right panel at 8.3, 10.2, and 14.9 mm per frame irregularity for the Hough transform.

## 4 Discussion

We presented a novel hybrid method for tracking muscle architecture from ultrasonography using a combination of a sequential (i.e., feature-point tracking) and non-sequential (i.e., Hough transform) method. Our results demonstrate that this approach can overcome the respective limitations of drift and irregularity of the individual methods, while tracking fascicle pennation angles and aponeurosis displacement of the tibialis anterior muscle during isometric contractions.

To start, the presented hybrid method was shown to mitigate the drift observed in two sequential optical flow estimation methods (i.e., a custom feature-point tracking approach and UltraTrack). It is evident from the two examples shown in Figure 4 that drift can lead to considerable deviations of the measured muscle architecture over time. These example trials were collected over twenty seconds of isometric contractions. However, ultrasonographic recordings of muscles during movement (such as walking or running on a treadmill) can involve trials of several minutes in duration. Tracking muscle architecture over such extended periods of time is likely to lead to further accumulation of tracking errors and excessive drift, thus rendering sequential methods unsuitable for longer ultrasound recordings. To the best of our knowledge, the presented hybrid method is the first muscle architecture analysis method that overcomes drift without the need for additional information (e.g., ground reaction forces) besides ultrasonography or manual input (e.g., as required for key-frame correction [5]). Our method thus opens new opportunities for tracking muscle architecture for extended ultrasound videos captured during movement.

Besides mitigating drift, our hybrid method was demonstrated to significantly reduce the irregularity (i.e., noise-induced fluctuations) of the aponeurosis displacements resulting from the non-sequential Hough-transform method. The examples shown in Figure 5 reveal clear irregularities that can result from using the Hough transform to determine aponeurosis displacements from estimated pennation angles. Such inconsistencies over time are primarily caused by the variability of noise in ultrasound images across different states of the muscle. Since B-mode ultrasound imaging is based on the acoustic impedance of tissues in the scanning region, muscle movement and contraction during scanning do not only produce visual changes in muscle architecture but also cause irregular alterations of the noise and attenuation characteristics. Such undesirable characteristics are sensitive to the density or anisotropy of the tissue and introduce blurred patterns of speckles or stripes in the ultrasound image, which can affect the result of feature extraction methods such as Hough transform. Although this inherent noise and consequent irregularity issue can be addressed by using e.g., different filtering techniques, such approaches can be arbitrary and lead to the loss of important information. To the best of our knowledge, our hybrid approach is the first muscle tracking method that overcomes irregularities based on sequential tracking data, and thus provides an objective means to address an inherent limitation of ultrasound imaging.

Muscle architecture studies have traditionally relied on the manual annotation of ultrasonographic images, but recent developments have allowed for the (semi-)automation of the muscle tracking process that requires limited (e.g., [5], [6], [21]) or no (e.g., [22], [23]) manual input. Full automation is of particular interest when analyzing large datasets in which manual initialization or annotation is unfeasible. The presented hybrid muscle tracking method allows for full-automatic detection of the aponeurosis locations and fascicle ROIs without the need for any user input. A major advantage of this approach is that it prevents the need for manually estimating the location of the fascicle insertion points in the aponeurosis if these are outside of the field of view (such as required in e.g., UltraTrack [5]), which thus eliminates experimenter bias and errors.

Our results suggest that the proposed hybrid method can reliably track tibialis anterior muscle architecture in most ultrasound videos for various participants. It should however be noted that, like other tracking methods, the performance of our algorithm depends on the choice of several hyperparameters, e.g., parameters related to the Frangi vesselness filter, aponeurosis search areas, and Hough transform (post-)processing options. In particular, the weight factor (w in Equation 1) plays a crucial role in controlling how much emphasis is put on the suggestion from feature-point tracking compared to that from Hough transform. Although our results confirm that the chosen value for w (i.e., 12.0) used in our study works robust for most videos, we found a small number of videos where adjustment of the weight factor can be considered. For example, two (outlier) trials shown in Figure 7 indicate that the fascicle pennation angle determined from the Hough transform did not follow the ankle moment pattern well, and consequently also dominated the aberrant pennation angle profile result from our hybrid method. In this case, increasing the weight factor to make our algorithm rely more heavily on the feature tracking algorithm can be considered. It is possible that such adjustments may also be required when our proposed algorithm is applied to a wider variety of ultrasonographic videos of muscle contractions, taken from different muscles and a range of ultrasound devices.

**Figure 7:**
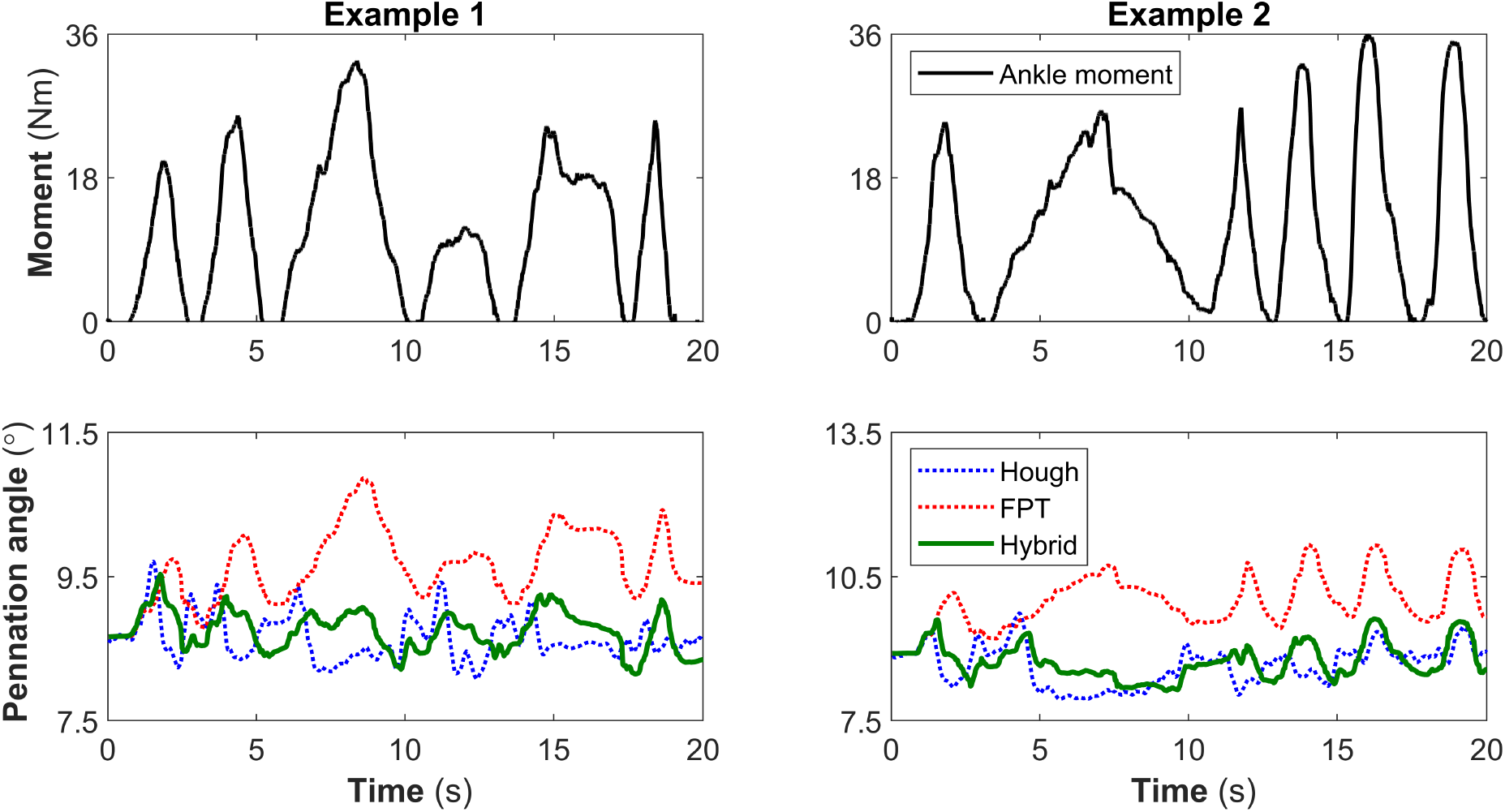
Fascicle pennation angle results for two example trials for which the tracking results of the proposed hybrid method (green solid line) were deemed unsatisfactory, primarily due to poor Hough-transform results (blue dotted line) that were not sufficiently corrected for by the feature-point tracking method (red dotted line). Both examples are for participant five at an ankle angle of 30° plantarflexion.

A common application for muscle architectural properties is the examination of the relationship between muscle structure and force production, or the direct estimation of muscle force given the state of the muscle. It is, however, well known that individual architectural properties do not correlate well with a muscle’s force output during different types of contraction. For example, although the pennation angle of muscle fascicles changes with the amount of force produced by the muscle, pennation angle by itself is known to be a poor predictor of a muscle’s force output [24], [25]. A combination of multiple architectural properties is thus likely required to gain a more comprehensive understanding of how a muscle’s overall architecture changes during force production. Our hybrid method provides a combination of multiple muscle architecture properties: fascicle pennation angle and length, and aponeurosis displacement, and thus allow for analyzing the intricate relationships between several muscle architectural properties and force output.

In this study we investigated the superficial part of the tibialis anterior muscle belly which has a relatively simple muscle architecture. As such, muscle fascicles were mainly straight and aligned in parallel, with a near-constant thickness between aponeuroses. This relatively simple structure allows for describing the muscle’s architecture using only a few properties and makes it possible to compare the performance of different tracking methods. However, a complication for evaluating different tracking methods is that a clear gold standard to provide ground truth data for muscle architectural properties does not exist. Even manual tracking is highly susceptible to human errors since, as can be seen from the result of tracking in Figure 5, the variation of the pennation angle is around 3° at most. Therefore, the comparison between tracking methods presented above can thus be insightful but cannot and should not be taken as the sole measure of accuracy. Furthermore, it should be noted that many human skeletal muscles comprise curved fascicles and aponeuroses, as well as non-uniformly aligned fascicles. Like most other existing muscle tracking methods, our hybrid method does not account for these more complicated muscle architectures. However, more sophisticated methods for quantifying fascicle curvature have been explored [26], [27], and could be implemented to complement the information provided by our hybrid method. Moreover, if aponeurosis curvature outside of the field of view is likely, ultrasound probes with a longer field of view or multiple ultrasound probes in series [28], or the application of alternative extrapolation methods (e.g., polynomial functions [8]), may be used instead of the linear extrapolation approach taken in this study. Depending on the architecture of the muscle of interest, we encourage future investigations to incorporate alternative sophisticated non-linear feature detection and tracking algorithms, in combination with our presented hybrid approach (i.e., combining a sequential and non-sequential method).

## 5 Conclusion

This paper demonstrates that using a combination of a feature-point tracking and a Hough transform method for measuring tibialis anterior muscle architecture from ultrasonography, can uniquely mitigate drift and irregularity limitations of the individual methods. The fully automatic nature of this hybrid method allows for the convenient analysis of large datasets of ultrasonographic videos. Moreover, automatic drift correction opens the door for tracking muscle architecture in long ultrasound recordings during common movements, such as walking, running, and jumping. Based on the results reported in this paper we anticipate that our hybrid method can be extended well to other human skeletal muscles, and we encourage future work to incorporate more advanced non-linear feature-detection and tracking algorithms to further enhance the accuracy and scope of available muscle architectural properties.

